# Effective Multiple Scattering Correction for Accurate Quantitative OCT

**DOI:** 10.1101/2025.02.28.640668

**Authors:** Cheng-Yu Lee, Xingde Li

## Abstract

We present an effective approach to compensate for multiple scattering effects in Quantitative Optical Coherence Tomography (qOCT) imaging, with the goal of accurately extracting tissue attenuation coefficients, which is crucial for precise clinical diagnosis. In clinical practice, especially for intra-operative imaging, an increased working distance is often necessary to avoid interference with surgical instruments and workflow. However, this increased working distance corresponds to an increased beam spot size, leading to more multiply scattered photons in the OCT signal and thereby underestimating the optical attenuation. To address this challenge, we investigated errors in attenuation coefficient quantification under different beam spot sizes. Monte Carlo simulations were employed to generate virtual OCT signals for two distinct beam spot sizes, enabling us to quantify the errors induced by multiple scattering across a range of true optical attenuation coefficients. Based on this analysis, we developed a compensation function to correct these errors. The proposed method was validated through experimental measurements using tissue-mimicking phantoms and demonstrated a significant improvement in the accuracy of attenuation coefficient quantification. Our results underscore the potential of this easy-to-implement technique to enhance the diagnostic reliability of qOCT, facilitating its broader application in clinical settings for accurate tissue characterization.

## 1. Introduction

Quantitative OCT (qOCT), an imaging technique based on the extraction of inherent optical properties of tissues from depth-dependent OCT signals, has demonstrated robust capability in tissue differentiation. In particular, attenuation-based qOCT, which harnesses the optical attenuation coefficients, has proven effective in detecting brain cancer with high sensitivity and specificity due to the distinct attenuation values between cancerous and noncancerous brain tissues. [1]. Typically, qOCT analysis assumes that the OCT signal is dominated by single-scattering photons, and the attenuation coefficient can be conveniently quantified using methods such as exponential/linear fitting [2], the robust frequency-domain algorithm (FD-method) [3], and depth-resolved methods [4-6]. However, the contribution of multiply scattered photons to the OCT signal becomes pronounced as depth and/or beam spot size increase, and the OCT signal appears to decay more slowly, leading to an underestimation of the optical attenuation coefficient using those methods. For many clinical applications, particularly for intra-operative imaging, a long working distance is needed to ensure sufficient space between the OCT imaging device and the tissue surface and avoid any interference of the OCT imaging probe with surgical tools. A long working distance corresponds to a large beam spot size, which makes depth-dependent OCT intensity signals more susceptible to multiply scattered photons, compromising the accuracy in attenuation quantification.

This study aims to analyze the impact of multiply scattering on attenuation-based qOCT accuracy and develop a practical and convenient method to remedy the impact of multiply scattering, with a goal of obtaining accurate optical attenuation. We began by selecting a beam spot size of 60 µm, corresponding to a working distance of 60 mm, suitable for intra-operative brain cancer imaging. OCT imaging was conducted with the beam implemented on five scattering phantoms, each with a distinct scattering-dominating optical attenuation coefficient, ranging from 1 mm^-1^ to 9 mm^-1^ to represent different tissue conditions. In parallel, we performed Monte Carlo (MC) simulations to generate virtual OCT signals from computer phantoms of the above optical attenuation coefficients (again ranging from 1 mm^−1^ to 9 mm^−1^). The simulations used the same optical properties and imaging beam configurations as the physical phantom experiments, allowing us to quantify the impact of multiply scattering on the attenuation coefficient characterization. By applying linear fitting to the logarithm of the OCT depth-dependent signals, we retrieved the attenuation coefficients from both the measured and simulated OCT signals. Analysis of the results confirmed that multiply scattering caused underestimation, as expected. To correct this, we developed a compensation scheme that restored the attenuation coefficient to its true value. To validate the effectiveness of this correction, we applied it to the measured data from the 60 µm beam spot size experiments. The corrected results showed a significant improvement in accuracy, closely matching the expected values based on the known optical properties of the phantoms. This confirmed that our method works well in compensating for the underestimated optical attenuation owing to multiple scattering and provides a more reliable quantitative assessment of the attenuation coefficient, demonstrating its potential for clinical applications.

## 2. Materials and Methods

### 2.1 OCT System and optical phantom

The swept-source OCT (SS-OCT) system used in this study has been described previously [7]. Briefly, our SS-OCT system employs a VCSEL laser operating with a 100 kHz A-line scanning rate, a central wavelength of 1310 nm, and a 3dB spectral bandwidth of approximately 90 nm (110 nm at 10 dB). The internal linear *k*-clock enables an imaging range up to 6 mm in air. The measured axial resolution is 9 µm in air, while the transverse resolution varies with different lens combinations, offering 20 µm and 60 µm options. Detection sensitivity at a 9.2 mW incident power on the sample is -115 dB, with a signal roll-off of -0.06 dB/mm. The B-frame image size is set at 2048 × 2048 pixels, representing a lateral × axial field of 2 mm × 6 mm in air. For this study, we fabricated optical phantoms of different optical attenuation coefficients. A reference phantom of 1.3 mm^−1^ scattering and 0.01 mm^−1^ absorption at 1310 nm (wavelength) consisting of 5% gelatin mixed with silicon dioxide (SiO2) nanospheres, averaging 180 ± 20 nm in diameter at a concentration of 50 mg/ml, was prepared. The optical properties of these silica nanoparticles have previously shown a strong correlation with the predictions of Mie theory, as confirmed in a prior calibration study [8]. All other tissue phantoms were constructed using SiliGlass (RTV silicone) embedded with titanium dioxide (TiO_2_) nanoparticles (SIGMA) as the scatterers, following a published protocol [9]. The TiO_2_ weight concentrations in these RTV silicone phantoms were 0.30%, 0.50%, 0.70%, and 0.90%, respectively, corresponding a predicted linear relationship of optical attenuation coefficient of 3 mm^−1^, 5 mm^−1^, 7 mm^−1^, 9 mm^−1^ [4]. OCT images were collected from each phantom with the light focused approximately at 300 µm beneath surface, from which optical attenuation coefficients were quantified for these phantoms of different scatter (TiO_2_ nanoparticle) concentrations. For beam spot sizes of 20 µm and 60 µm in diameter, this depth remains within one Rayleigh length and has been empirically chosen in our imaging practice, which balances the imaging signal and contrast degradation along depth.

### 2.2. Monte Carlo simulations for simulated OCT data

Assessing the accuracy of qOCT and the impact of multiply scattering on the quantified optical attenuation requires OCT imaging data from robust tissue mimicking phantoms. However, in the fabrication of the above phantoms (or any tissue phantoms in general), issues such as nonuniform distribution of the scatters due to clustering or incomplete mixing and the formation of air bubbles will introduce errors in attenuation coefficient calculations. Due to these limitations, we opted for Monte Carlo simulations, which offer precise control over scattering parameters. This approach allows for a systematic analysis of multiple scattering effects on OCT measurements and attenuation quantification. To simulate OCT intensity versus depth for various optical attenuation (scattering) properties, we adopted the well-established MCEO-OCT framework, an adaptation of the MCML algorithm designed for modeling light-tissue interactions [10, 11]. To make the simulations more consistent with realistic settings, we modified the framework to incorporate the Gaussian beam focusing effects [12-14].

Figure 1 illustrates the schematic of a Gaussian incident beam. Incorporating a Gaussian beam model into a standard MC simulation involves changing the initial distribution of the photons before their first interaction with tissue. The beam radius at depth *z* can be calculated from the Gaussian beam formula:

**Figure 1.**
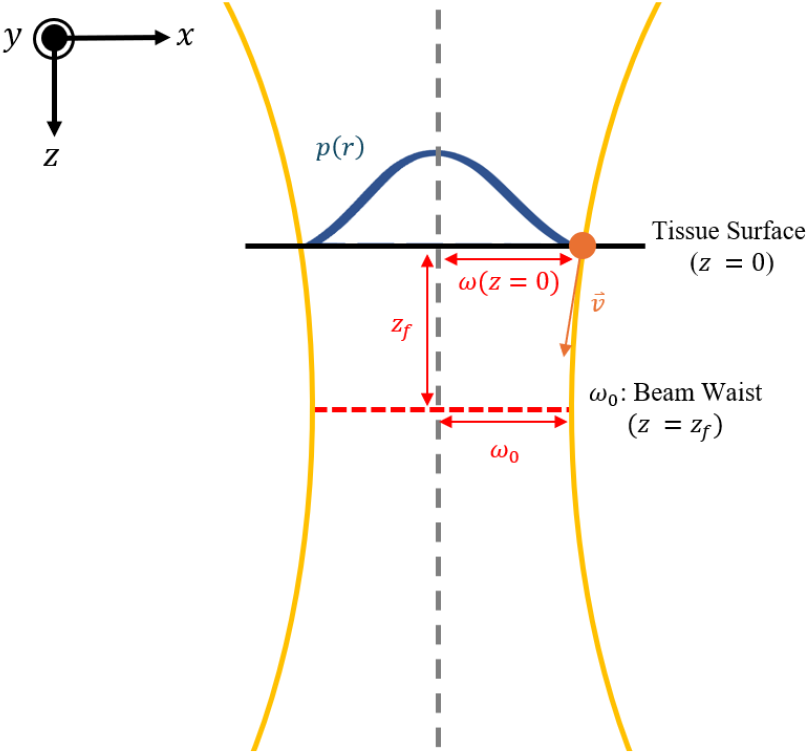
Schematic of the Gaussian profile of the incident OCT beam. *p*(*r*): probability density function of the incident photon;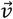: direction unit vector of a given incident photon; *z*_*f*_: the focusing position; *ω*_0_: beam waist, 2*ω*_0_ is the beam spot size, *ω*(*z* = 0): beam radius at the tissue surface.

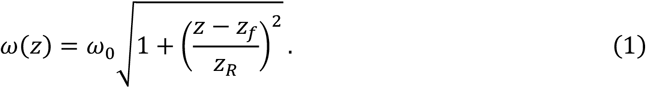

Therefore, the incident beam radius at the tissue surface (*z* = 0) is given by:

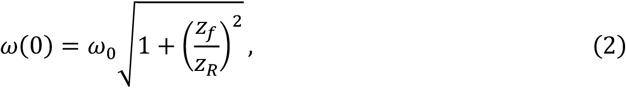

where, *ω*_0_ is the beam waist radius, 2*ω*_0_ is the beam spot size, *z*_*R*_ is the Rayleigh length, and *z*_*f*_ is the focus position. The probability density function *p*(*r*) of the photon incident location was defined by the intensity distribution of the Gaussian beam shown below [15]

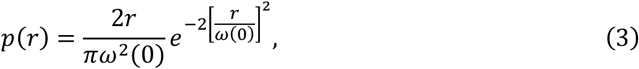

where *r* is the lateral distance from the beam center at the tissue surface. The direction unit vector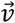of a photon incident at (*x, y*) on the tissue surface, is given by

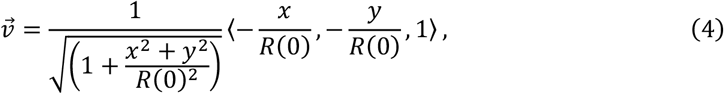

where *R*(0) is the curvature of the beam profile at tissue surface, which can be expressed as

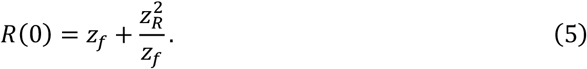

The photon keeps traveling in the direction specified by Eq. (3) until its first scattering event, after which it is tracked using the standard MCML. The remaining simulation process, including the use of the Henyey-Greenstein phase function to update scattering angles, photon weight adjustments using the Russian Roulette method, and photon tracking, followed the established framework [10, 11].

The depth-dependent OCT interference signal from MC simulations was generated by tracking photon interactions as they traveled through the simulated tissue medium. Each photon packet underwent scattering and absorption events based on predefined optical properties, including the scattering coefficient, absorption coefficient, and anisotropy factor. Each backscattered photon was then categorized into a *z* bin based on the path length which it traveled with a coherence gating [12]. The frequency of the *z* bins eventually represent the depth-dependent backscattered signal, and the OCT interference signal is proportional to the square root of the backscattered signal. To investigate the impact of beam size on attenuation quantification (due to multiple scattering), MC simulations were conducted with two beam spot sizes: 20 µm and 60 µm. The 20 µm spot, with a working distance of approximately 20 mm, has a smaller illuminated area, and it is expected to result in reduced multiple scattering effects and thus lower quantification error. In contrast, the 60 µm spot, with a working distance of around 60 mm (roughly three times larger than the former), covers a larger area, which is expected to lead to increased susceptibility to multiple scattering and quantification errors. Comparing these two beams in the simulations allowed us to analyze how multiple scattering influences attenuation quantification under differing spot sizes and working distances. For our simulation parameters, scattering coefficients ranged from 1 mm^−1^ to 9 mm^−1^, with a fixed absorption coefficient of 0.01 mm^−1^. The refractive index was set to 1.4, and the anisotropy factor to 0.75, to closely match experimental conditions. To reduce speckle noise, each simulation was repeated 30 times and then averaged for data smoothing.

### 2.3. Quantification of the attenuation coefficient

We compared A-line signals experimentally measured from the physical tissue phantoms with the corresponding ones generated by MC simulations for optical attenuation coefficients ranging from 1 mm^−1^ to 9 mm^−1^. The linear fitting method [2, 8] was used to calculate attenuation coefficients for our two selected beam sizes: 20 µm and 60 µm. To minimize intensity variations caused by the Gaussian beam profile effect at 300 µm depth, which deviates from the single-scattering assumption, the OCT signals before fitting [8]. For measured OCT data, we divided the signal by that of a reference phantom to normalize the intensity. For simulated OCT data, we normalized the signal by the phantom with the lowest simulated attenuation coefficient (1 mm^−1^) to reduce the Gaussian profile effect. After taking the logarithm of the depth-dependent OCT signal, we performed linear fitting over a 200 µm region starting from 100 µm beneath surface to avoid possible errors from strong surface reflections. This approach enabled a systematic comparison of attenuation values across different beam sizes and scattering conditions, providing insights into the impact of beam size on attenuation quantification.

### 2.4. Compensation method

To address the challenges posed by multiple scattering effects (particularly when using large beam spot sizes), which lead to an underestimation of the attenuation coefficient, a strategic approach was implemented through the development of a meticulously function which compensates for the underestimate of the quantified attenuation coefficient *μ*_*att*_ [16]. Here we explore an easy-to-implement method. As described in Section 2.2, we adopt the results from MC simulated OCT signals to obtain the compensation function, by fitting the attenuation coefficient *μ*_*att*_ quantified from the MC simulated depth-dependent OCT signals to the theoretical values *μ*_*true*_, the ones used in the MC simulations) with a third-order polynomial function:

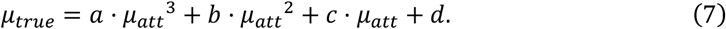

The simplified model is implemented for its balance of flexibility and simplicity, which effectively captures the non-linear relationship between underestimated and true attenuation coefficients.

## 3. Results

### 3.1. MC simulation results and measured OCT data

To serve as a basis for quantified attenuation coefficient compensation, MC simulation results should closely resemble the OCT data measured correspondingly. In order to clearly demonstrate the validity MC simulation results, we compare them with the measured OCT data.

Figure 2 displays the simulated results for the scattering coefficient of 1 mm^−1^, 3 mm^−1^, 5 mm^−1^, 7 mm^−1^, and 9 mm^−1^ at 20 µm and 60 µm beam spot size. While the simulated and experimentally measured A-line profiles are not perfectly identical, they exhibit a high degree of similarity, fulfilling the aforementioned requirement.

**Figure 2.**
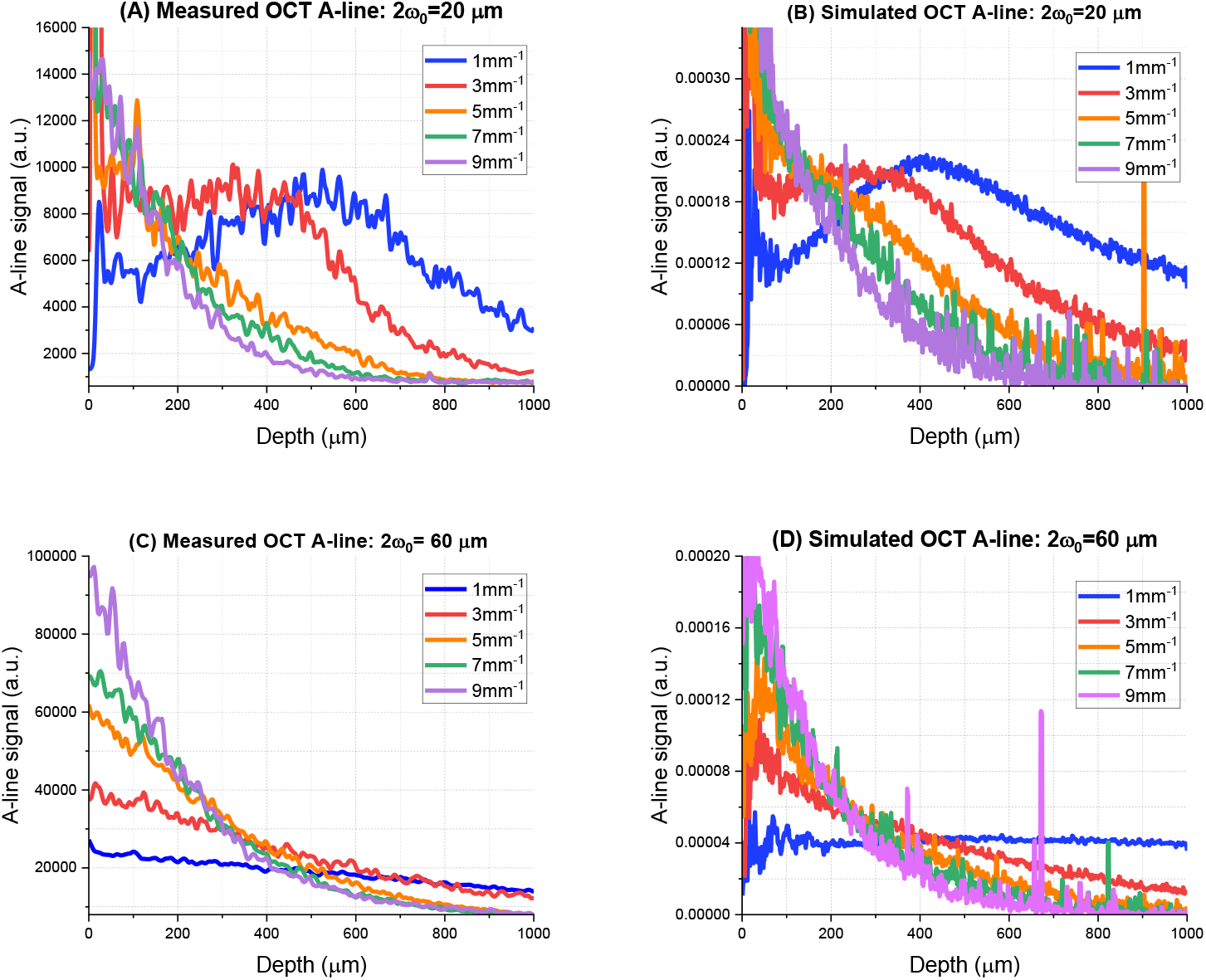
Comparison between the simulated result and the measured OCT data, with the beam focused ∼214 µm (300 µm in air) beneath the tissue: (A) 20 µm beam spot size, measured OCT A-line. (B) 20 µm beam spot size, simulation OCT A-line from Monte Carlo Simulation. (C) 60 µm beam spot size, measured OCT A-line. (D) 60 µm beam spot size, simulation OCT A-line from Monte Carlo Simulation.

### 3.2. Results of attenuation quantification

We quantitative compared attenuation coefficients obtained from experimental and simulated data, enabling an evaluation of our quantification method’s accuracy across varying optical attenuation values and beam sizes.

Figure 3 presents the attenuation coefficients quantified from the simulated and measured OCT A-line data. Symbols in red color represent data points obtained from MC simulated and measured A-line signals when using a 20 µm beam spot size, while the symbols in blue color represent data points when using a 60 µm beam spot size. Squares represent results from MC simulated data and triangles represent results from measured OCT data. The dashed black line represents the ideal correlation between the quantified attenuation and the true attenuation. As shown in Fig. 3, the measured attenuation coefficients closely align with the MC simulation results, validating the accuracy of our experimental approach. It is also evident that the 60 µm beam size tends to underestimate the attenuation coefficient more severely compared to the 20 µm beam size, particularly at higher true attenuation values. This discrepancy highlights the importance of considering beam size in quantitative OCT measurements to ensure accurate attenuation coefficient estimation.

**Figure 3.**
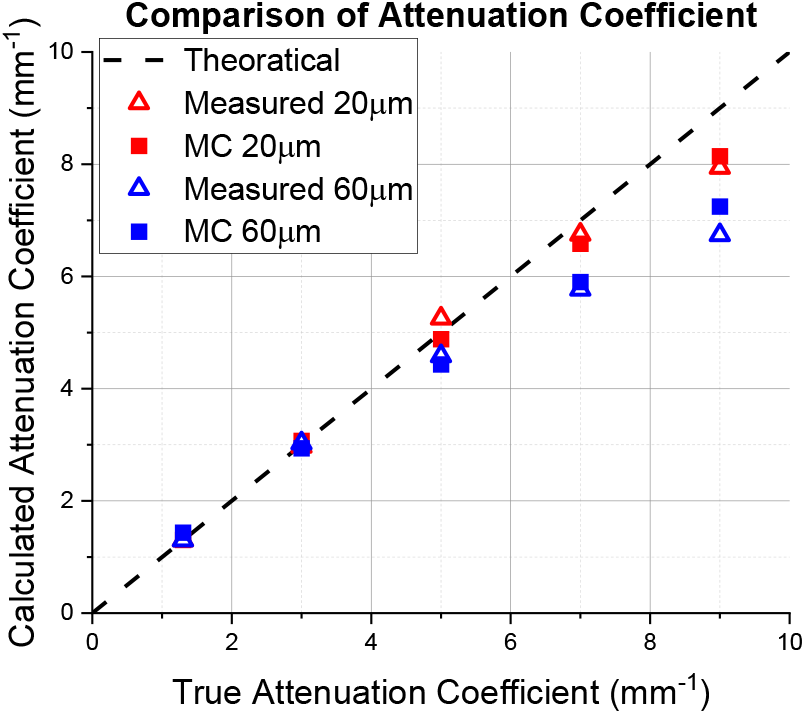
Comparative analysis of measured and simulated OCT Data. Dashed black line: ideal correlation of calculated attenuation and the true attenuation. Red triangles: measured qOCT results with a 20 µm beam spot size. Red squares: the MC simulation results at a 20 µm beam spot size. Blue triangles: measured qOCT results at a 60 µm beam spot size. Blue squares: the MC simulation results at a 60 µm beam spot size.

### 3.3. Compensation function for attenuation quantification

In the clinically relevant threshold range of attenuation coefficients (∼4.5 mm^−1^) for brain cancer detection, the attenuation coefficients measured with the 20 µm beam spot size closely matched the theoretical values, indicating that compensation was unnecessary. In contrast, measurements with the 60 µm beam spot size exhibited significant underestimation due to multiple scattering, necessitating compensation.

To address this, the compensation function developed in Section 2.4 was applied to OCT measurements taken with a 60 µm beam spot size to correct for attenuation coefficient underestimations caused by multiple scattering. By fitting the results obtained from the simulated OCT data to this polynomial function, we found the coefficients in Eq. 7, and the final compensation function becomes

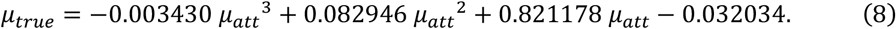

The effectiveness of the compensation function was validated by comparing the corrected attenuation coefficients quantified from OCT imaging data with a 60 µm beam spot size with true attenuation coefficients.

As shown in Fig. 4, the corrected attenuation coefficients (red dots) closely align with true values, demonstrating the effectiveness of the compensation function in reducing the underestimation errors caused by multiple scattering. This confirms the reliability of our compensation approach, particularly for larger beam spot sizes where multiple scattering effects are more pronounced.

**Figure 4.**
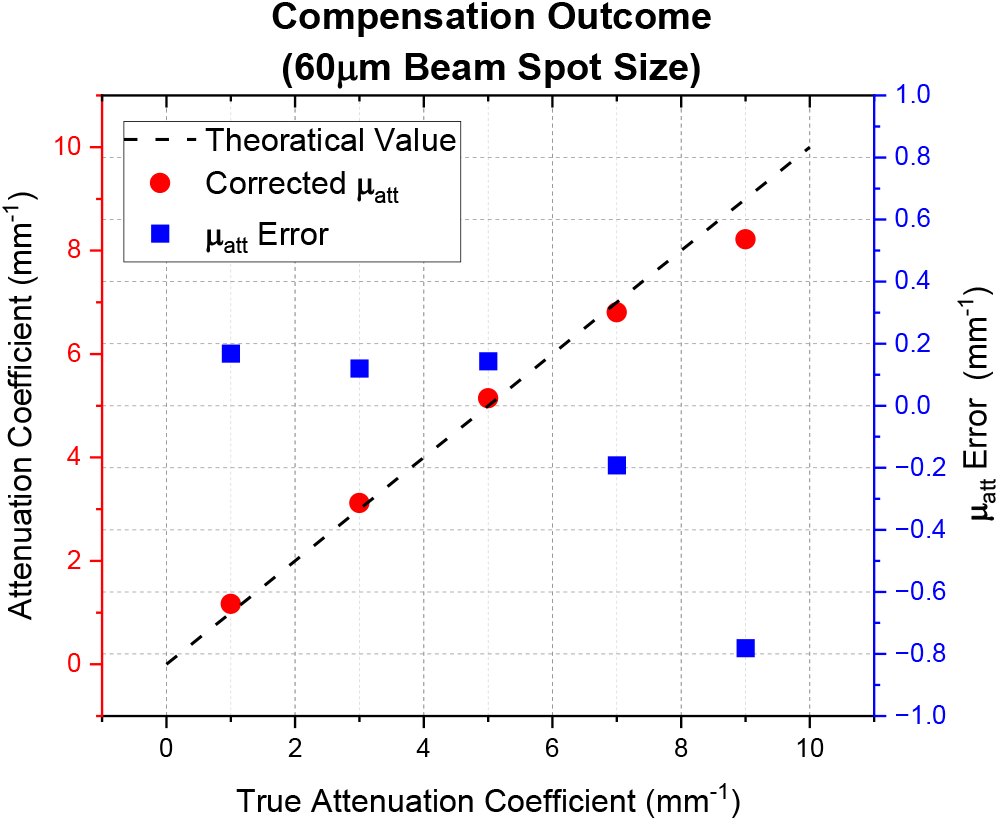
Compensation outcome for attenuation quantification at a 60 µm beam spot size. Dashed black line: ideal correlation of calculated attenuation and the true attenuation. Red dots: corrected attenuation (i.e., quantified attenuation coefficients from OCT measurements after compensation). Blue rectangles: attenuation error (i.e., the difference between quantified attenuation from OCT imaging data after compensation and the true attenuation).

However, at high attenuation (7 mm^-1^ and higher), the measured attenuation coefficients with the 60 µm beam spot remained lower than the true values, leading to a residual underestimation even after compensation. Considering the threshold for cancer detection is around 4.5 mm^-1^, the remaining discrepancy would not categorically change the diagnostic conclusion. For measured attenuation around the cancer detection threshold, e.g., in the range of 4.0–5.0 mm^−1^, the discrepancy in attenuation measurements after compensation was smaller than 0.15 mm^−1^. According to previous studies, this level of error is expected to reduce the sensitivity or specificity of brain cancer detection by less than 1.3%, ensuring that the compensation function can effectively restore quantitative accuracy for intraoperative applications [17].

### 3.4. Validation on phantom imaging with compensated qOCT

To validate the efficacy of the compensation function derived from MC simulations in correcting the underestimated attenuation coefficients due to multiple scattering, we constructed a phantom with two distinct attenuation levels.

Figure 5 (A) displays a photo of the test phantom, featuring two compartments with low attenuation (left: with ∼0.30% TiO_2_ weight concentration) and high (right: with ∼0.60% of TiO_2_ weight concentration). The objective of such settings was to loosely mimic human white matter imaged intraoperatively, with the low attenuation area representing cancerous tissue, the high attenuation area representing noncancerous tissue and the boundary of the two areas as an infiltration zone. Data collection was carried out using two different beam spot sizes (20 µm and 60 µm) and the region of interest (ROI) was set at the boundary between the two compartments, so that assessment of the capability of attenuation maps quantified with a 60 µm beam spot size to distinguish the two regions can be made (the robustness of attenuation maps quantified with a 20 µm beam spot size has been validated in earlier studies [1, 18]). Attenuation quantification was conducted using the LF method. This process generated a color-coded 2D attenuation map with red, green and yellow denoting cancerous, noncancerous and infiltrative regions for better visualization [1].

**Figure 5.**
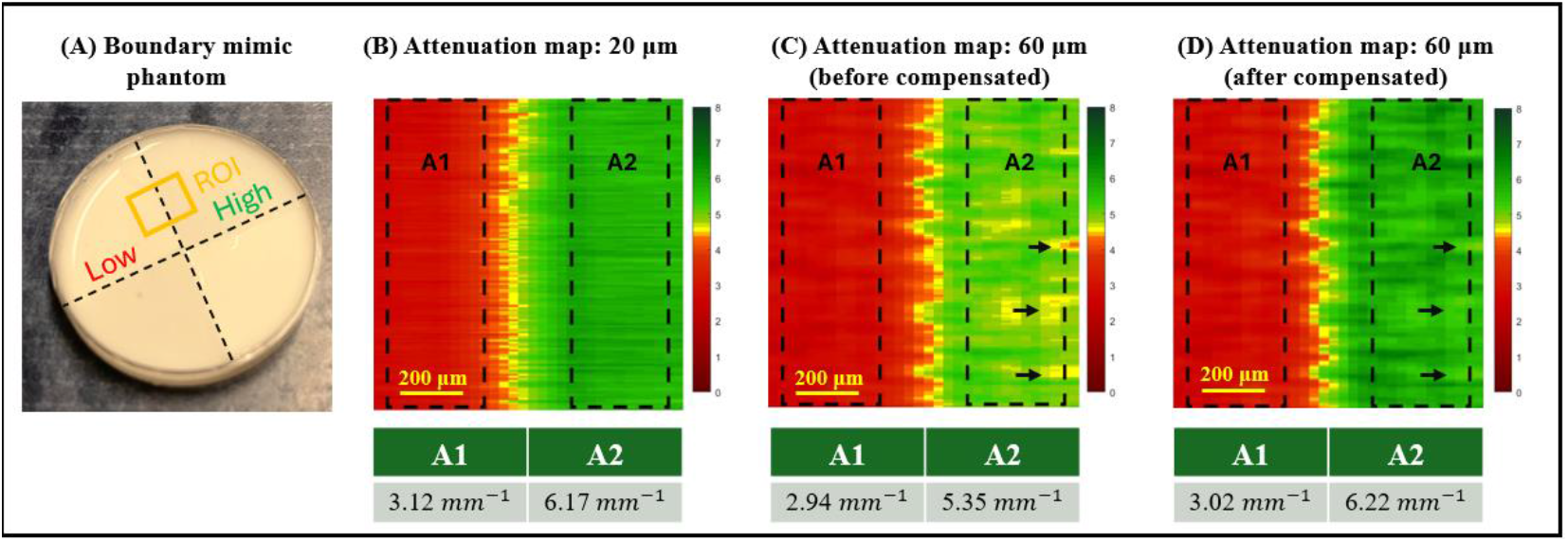
Illustration of color-coded attenuation maps. (A) Photo of the boundary mimic phantom. (B) Attenuation map obtained using a 20 µm beam spot size. (C) Attenuation map obtained using a 60 µm beam spot size before compensation. The arrows highlight representative areas where attenuation was underestimated. (D) Attenuation map obtained using a 60 µm beam spot size after applying the compensation function (Eq. 8). The arrows, corresponding to the same locations as in (C), indicate the restoration of attenuation after compensation, as reflected by the color changes. Note that the attenuation coefficients shown at the bottom of (B-D) represent the average values within the respective boxed regions.

Figure 5 (B) shows the color-coded attenuation map quantified from volumetric OCT image data acquired with a 20 µm beam spot size across the boundary, clearly delineating the low and high attenuation areas. The average attenuation coefficient in area A1 was measured as 3.12 mm^−1^ and in area A2 as 6.17 mm^−1^, both slightly higher than the theoretical values (3 mm^−1^ and 6 mm^−1^, respectively). This deviation may be attributed to measurement inaccuracies introduced during the phantom fabrication process. Figure 5 (C) illustrates the color-coded attenuation map quantified from volumetric OCT image data acquired with a 60 µm beam spot size across the boundary. Without compensation, the map reveals localized underestimation of attenuation coefficients, particularly in regions near the 4.5 mm^−1^ threshold, leading to non-uniformity in the color-coded 2D attenuation map (as indicated by the arrows in the Figure 5(C)). Additionally, due to this underestimation, the boundary representing the infiltration zone is slightly shifted to the right. Figure 5(D) presents the compensated attenuation map, which exhibits similar features to Figure 5(B). Notably, the attenuation values at locations marked by arrows, which previously indicated underestimation in Figure 5(C), are effectively restored after compensation.

## 4. Discussion and Conclusion

In this study, we developed and validated a method to correct the underestimation of attenuation coefficients in qOCT measurements when implementing larger beam spot sizes, by introducing a compensation function that effectively addresses the underestimated attenuation coefficients caused by multiple scattering. This compensation method enables the use of larger beam spots and thus extended working distances are critical for intra-operative imaging applications.

For complex or multi-layered samples, a depth-resolved analyses of the attenuation coefficients could be used [4, 6]. However, multiple scattering again impacts the accuracy of attenuation quantification, leading to an overall underestimation of attenuation coefficients. Future research should further explore the actual proportions of multiple scattering and develop methods to mitigate these effects to align the signals more closely with a single scattering model.

In this study, by leveraging MC simulated OCT signals and comparing them with measured OCT signals from phantoms with the same optical properties, we gained better understanding of the factors affecting the accuracy of attenuation quantification under different beam spot sizes. This approach not only enhances the reliability of qOCT measurements but also facilitates the use of larger beam spot sizes with extended working distances, making it more suitable for intra-operative imaging applications. The findings lay a foundation for further research to optimize attenuation corrections in clinical settings, ultimately improving the precision of optical imaging in medical diagnostics.

